## Supplemental Files for "Generation of a bloodstream form *Trypanosoma brucei* double glycosyltransferase null mutant competent in receptor-mediated endocytosis of transferrin"

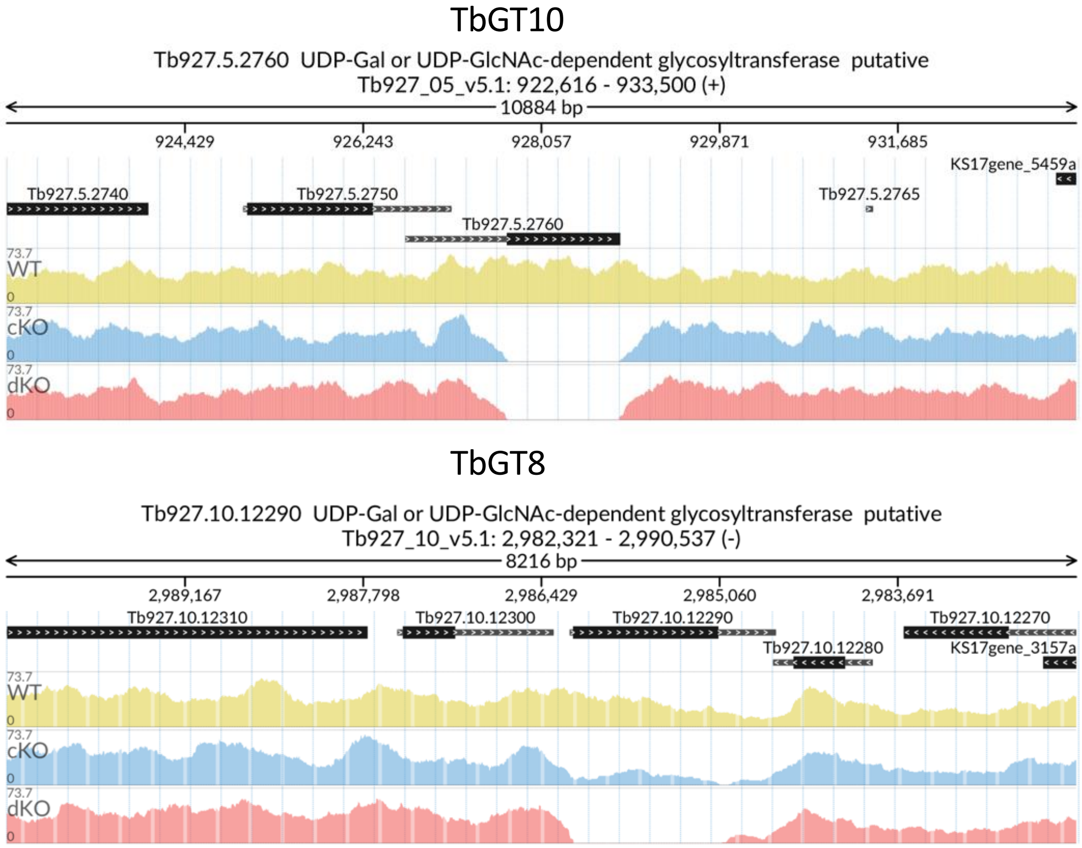


**S1 Fig. Genome sequencing validates knockout of both *TbGT8* and *TbGT10* in the *TbGT10*$^{-/-}$/*TbGT8*$^{-/-}$ double null mutant.** Genomic DNA harvested from WT, *TbGT10*$^{-/-}$/*TbGT8*$^{Flox/-}$ conditional null mutant (cKO) and *TbGT10*$^{-/-}$/*TbGT8*$^{-/-}$ double null mutant (dKO) cells was subjected to whole genome sequencing (30x coverage, paired end reads) and aligned to the *Trypanosoma brucei brucei* TREU927 reference genome. The plot shows mapped reads at the target genomic loci for *TbGT10* (Tb927.5.2760) and *TbGT8* (Tb927.10.12990).

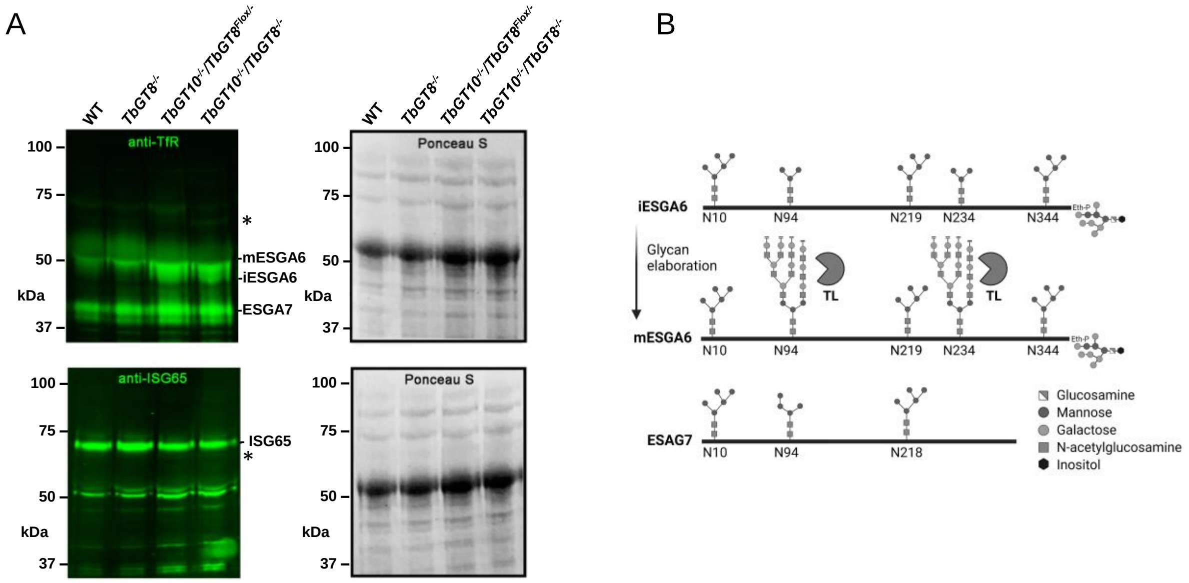


**S2 Fig. The RCA reactive glycoprotein band expressed in *TbGT10$^{-/-}$/TbGT8$^{-/-}$* double null mutants does not correspond to the transferrin receptor or invariant surface glycoprotein 65.** Cell ghosts were purified from soluble VSG (sVSG) by osmotic lysis. sVSG from WT, *TbGT8$^{-/-}$* null mutant, *TbGT10$^{-/-}$/TbGT8$^{Flox/-}$* conditional null mutant and *TbGT10$^{-/-}$/TbGT8$^{-/-}$* double null mutant cells were resolved by SDS-PAGE and transferred to nitrocellulose. Western blotting (green) with anti-transferrin receptor (TfR: upper panels) detecting both expression site associated genes (ESAG) 6 and 7 and anti-invariant surface glycoprotein 65 (ISG65: lower panels). Equal loading and transfer are demonstrated by Ponceau S staining (right). The region corresponding to the RCA reactive band is indicated (*) on the right and the molecular weight markers shown on the left. Products corresponding to mature (m)ESAG6, immature (i)ESAG6 and ESAG7 are indicated on the right. B. A schematic displaying the *N*-linked glycosylation states of immature and mature ESGA6 and ESAG7. Mature ESAG6 is reactive with tomato lectin (TL).

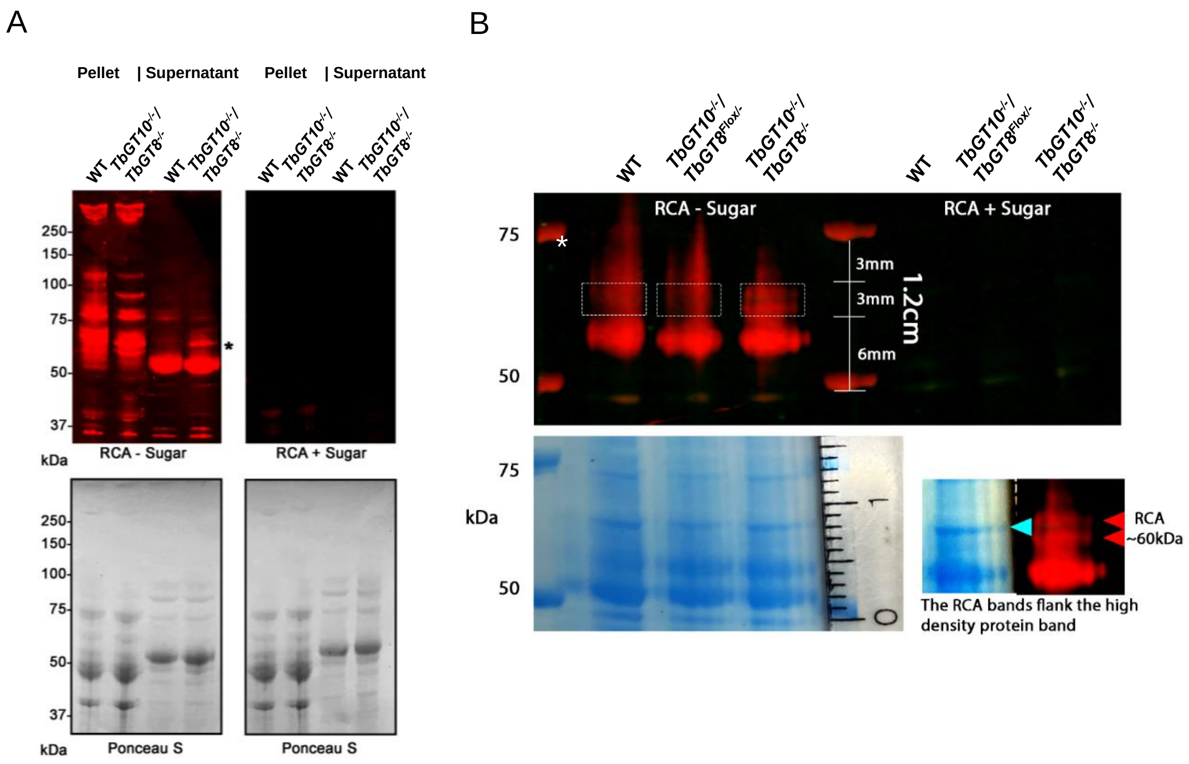


**S3 Fig Protein identification of glycoproteins exhibiting increased RCA reactivity in *TbGT10$^{-/-}$/TbGT8$^{-/-}$* double null mutants** A. Cell ghosts (pellet) were purified from VSG fraction (supernatant) by osmotic lysis from WT and *TbGT10$^{-/-}$/TbGT8$^{-/-}$* double null mutant cells. Pellet and supernatant fractions were resolved by SDS-PAGE and transferred to nitrocellulose in duplicate. Membranes were incubated with biotinylated RCA (red) without (RCA – Sugar: left) or with (RCA + Sugar: right) pre-incubation with 30 mg/ml galactose and lactose. Equal loading and sample transfer are demonstrated by Ponceau S staining. B. VSG (supernatant) fractions purified from WT, *TbGT10$^{-/-}$/TbGT8$^{Flox/-}$* conditional null mutant and *TbGT10$^{-/-}$/TbGT8$^{-/-}$* double null mutant mutants were depleted with anti-VSG221 conjugated agarose beads to remove VSG221 and non-bound material was incubated with RCA lectin agarose beads to purify RCA reactive glycoproteins. RCA agarose bound material was eluted by heating in SDS-sample buffer and resolved in triplicate lanes to enable RCA lectin blotting (upper panel) with RCA pre-incubated with 30 mg/mL galactose and lactose (RCA + Sugar) or not (RCA -Sugar) to enable Coomassie staining (lower panel). Gel slices from each sample corresponding to the RCA doublet were excised and submitted for proteomic analysis. The identified proteins are shown in Table 1.

**>ESAG2 Tb927.11.14620**
MEKLGILLLATTTFVVGTTTENKENLRNQQEFNQLCRILRLVEGDPGQVVKEPPNLGILIEVLQKMVKATFVKESSYQE
AVDAREKWKSTDSTVETKNIMDKQRADDLVRYKITHEKLKKLLAKAKMLVSRIKEERYLAYVSRNAALEKMAKVVYGS
SAEKMFRNETEFEKFLANSQGTIVGKTAKETCGMSNDKMKNVSFAGYSLVGDFFCLCVGENNNATLCDKSFSGPR
EEHKWTKVMFEDKLVDFAAGWFKIRGLCYVNATSGIKDVVTPENISTEIVAFNKMLGRQRDDVQIYRREAESYRQYIL
GRVEQVGFTDSAICTGEHQHMCVNYAQCLVSNKGIPWQLKLMEAEEDLEDVEKHNMEVHVLIGRLDKLIEKIKKAA
EQLSQEMEGIDVAVEKVQLLEEEIEENETTGKSEATAGEFNDNAQPEEGSLDKCEYWGMFSILFGVVAFSLV

| No. | Sequence | Position | Score | Prediction |
|---|---|---|---|---|
| 1 | YGSSAEKMFR[N]ETEFEKFLAN | 164 | 0.814613 | Complex/paucimannose |
| 2 | TCGMSNDKMK[N]VSFAGYSLVG | 197 | 0.151820 | Oligomannose |
| 3 | FFCLCVGENN[N]ATLCDKSFSG | 219 | 0.841290 | Complex/paucimannose |
| 4 | WFKIRGLCYV[N]ATSGIKDVVT | 262 | 0.063577 | Oligomannose |
| 5 | SGIKDVVTPE[N]ISTEIVAFNK | 275 | 0.859628 | Complex/paucimannose |
| 6 | VQLLEEEIEE[N]ETTGKSEATA | 412 | 0.707136 | Complex/paucimannose |

**>CBP1B Tb927.10.1040**
MMLCHTSLFLISLLLLLQSASALQAIRRPTLRTTGSGWEPCDPGVNQWSGYFDIPGEQSDKHYFYWAFGPRDGNP
NAPVLLWMTGGPGCSSMFALLAENGPCLMNETTGDIYNNTYSWNNHAYVIYIDQPAGVGFSYADKADYDKNEAEV
SEDMYNFLQAFFGEHEDLRENDFFVVGESYGGHFAPATAYRINQGNKKGEGIYIPLAGLAVGNGLTDPYTQYASYPR
LAWDWCKEVLGYSCISRETYDSMNSMVPACQSNISACDADNSSSADSYCEMAGAACSGFVSDFLLTGINVYDIRK
TCDGPLCYNTTGIDNFMNREDVQRSLGVDPMTWQACNMEVNLMFDIDWFKNFNYTISGLLEDGVRVMIYAGDM
DFICNWIGNKEWTLALQWSGSEEFVKAPDTPFSSIDGSAAGLVRSVSSNTSSMHFSFVQVYRAGHMVPMDQPAA
ASTIIEKFMRNEPLS

| No. | Sequence | Position | Score | Prediction |
|---|---|---|---|---|
| 1 | LLAENGPCLM[N]ETTGDIYNNT | 104 | 0.926133 | Complex/paucimannose |
| 2 | LMNETTGDIY[N]NTYSWNNHAY | 112 | 0.898209 | Complex/paucimannose |
| 3 | MNSMVPACQS[N]ISACDADNSS | 259 | 0.751451 | Complex/paucimannose |
| 4 | QSNISACDAD[N]SSSADSYCEM | 267 | 0.922695 | Complex/paucimannose |
| 5 | RKTCDGPLCY[N]TTGIDNFMNR | 310 | 0.452993 | Oligomannose |
| 6 | MFDIDWFKNF[N]YTISGLLEDG | 354 | 0.846613 | Complex/paucimannose |
| 7 | AAGLVRSVSS[N]TSSMHFSFVQ | 423 | 0.086370 | Oligomannose |

**S4 Fig *N*-linked glycan prediction analysis of ESAG2 and CBP1B.** The amino acid sequences from each protein were analysed using the N-glycosylation site prediction software [27]. Complex/paucimannose (red) and Oligomannose (green) glycosylation sites are indicated in the tables alongside their prediction scores and on the amino acid sequences. Signal peptides predicted by SignalP analysis are represented on the amino acid sequences in pink.

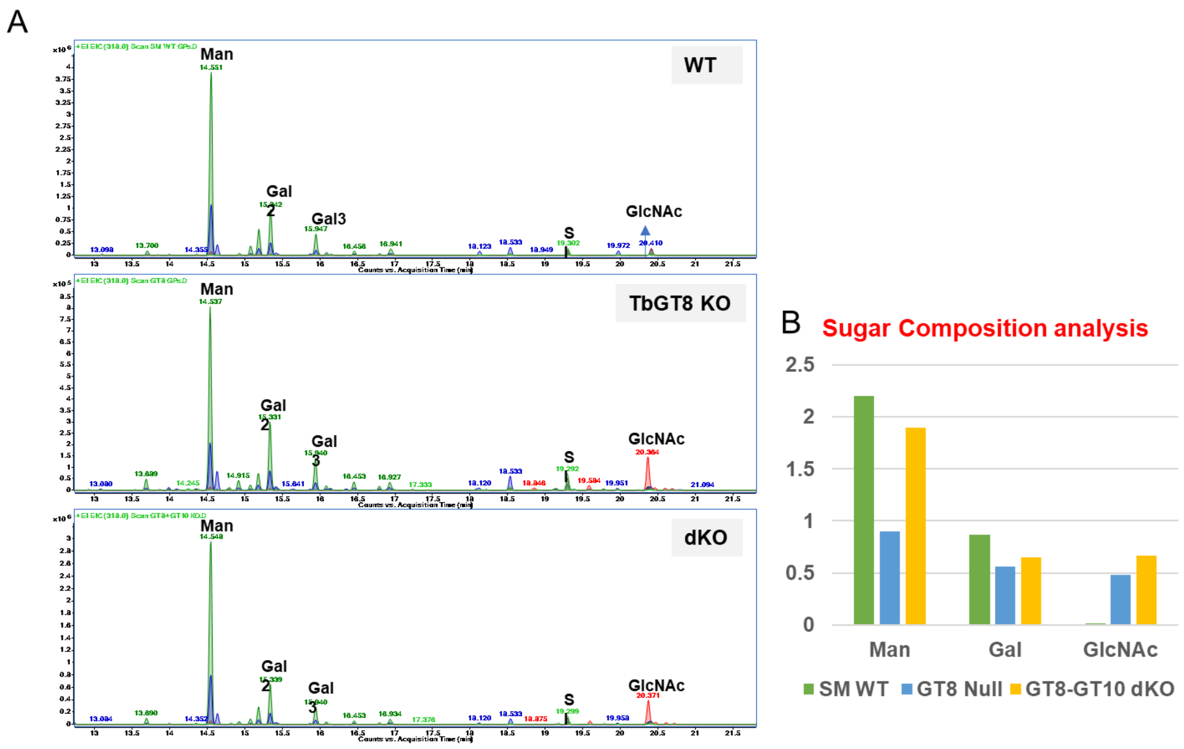


**S5 Fig GC-MS extracted ion chromatograms of wild type (upper) *TbGT8*$^{-/-}$ null mutant (middle) and *TbGT10*$^{-/-}$/*TbGT8*$^{-/-}$ double null mutant (lower) monosaccharide composition analysis.** A. The extracted *N*-glycopeptides were subjected to methanolysis and trimethylsilylation and the obtained methyl glycosides were analysed by GC-MS. 1 nmole of scyllo-inositol was used as an internal standard. B. Abundance of different monosaccharides plotted as a bar graph. Man: mannose, Gal: Galactose, GlcNAc: *N*-acetylglucosamine

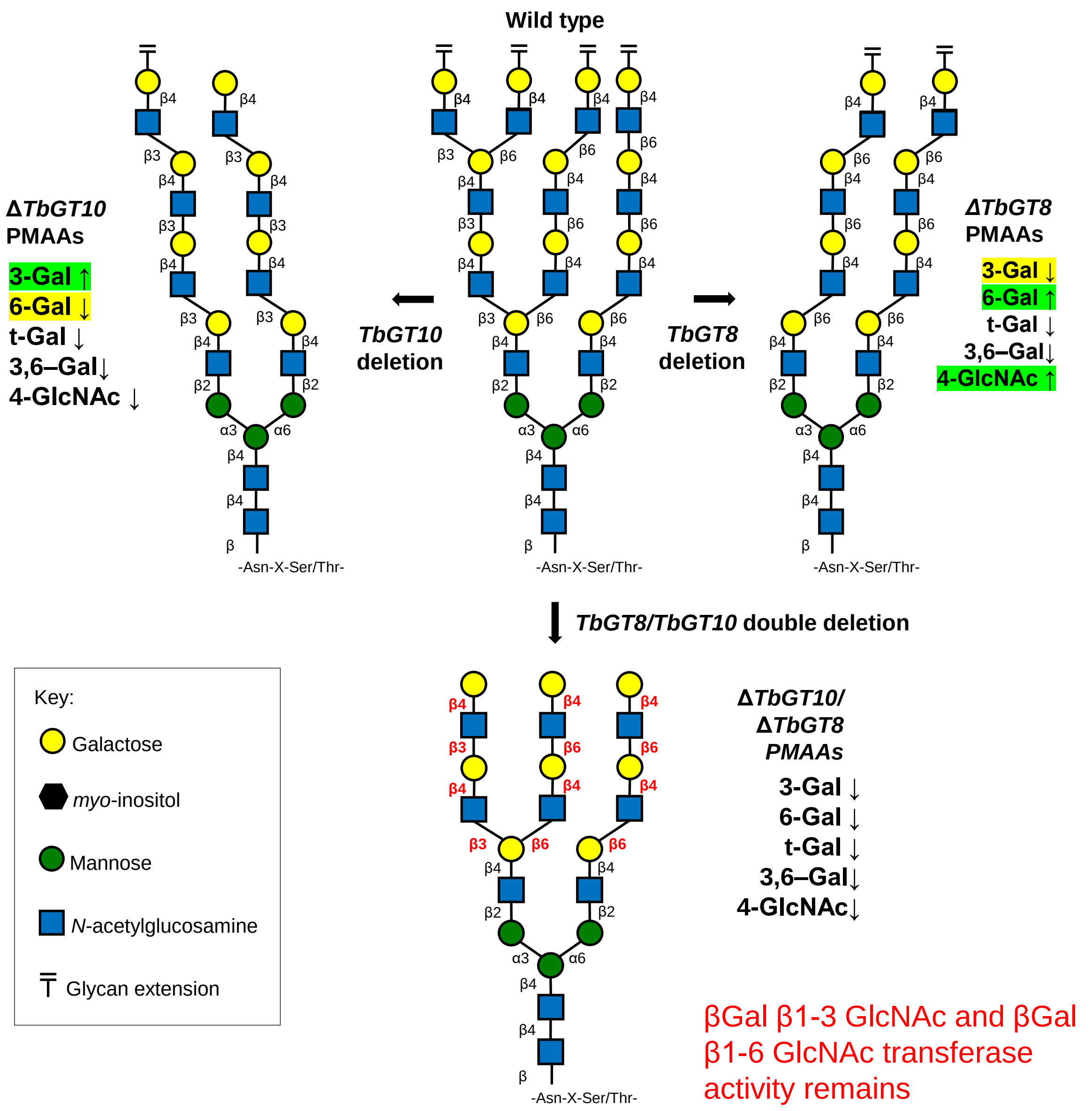


**S6 Fig Consensus glycoconjugate structures in TbGT mutant *T. brucei*.** Representative schematic of the complex *N*-glycan structures synthesised in cells depleted of TbGT10, TbGT8 or both (double deletion), based on permethylation linkage analysis.

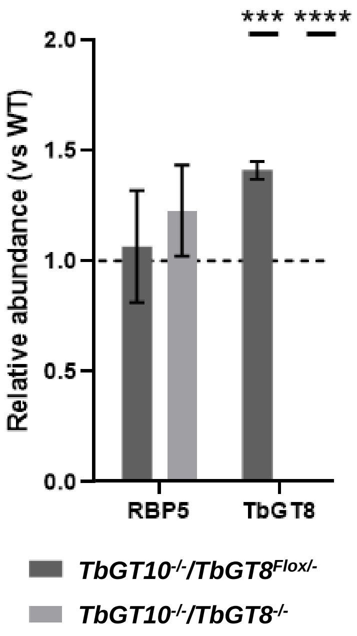


**S7 Fig: RBP5 expression is unchanged in *TbGT10$^{-/-}$/TbGT8$^{Flox/-}$* conditional null mutants and *TbGT10$^{-/-}$/TbGT8$^{-/-}$* double null mutants.** qRT-PCR analysis was performed to investigate *RBP5* and *TbGT8* transcript abundance in *TbGT10$^{-/-}$/TbGT8$^{Flox/-}$* conditional null mutant and *TbGT10$^{-/-}$/TbGT8$^{-/-}$* double null mutant cells, relative to WT (dashed line). Data are means ± SD (n=3 biological replicates with 3 technical replicates per n) ***P <0.0005 ****P <0.000005 as determined by unpaired T-test as compared with WT RNA expression levels

| Application | Oligo no. | Sequence | Amplicon |
|---|---|---|---|
| GT8 primers for pSY45_pSD66 flanking | SMD349 | aacgacggccagtgaattcgagctcCTTGCATCGCGACTGTTTTTCTTC | TbGT8 5’ homologous flank |
| | SMD350 | gcttctgcacttGCTGGGGTCCTTTCTTTCCTTG | |
| | SMD351 | aaaggaccccagcAAGTGCAGAAGCTTATAACTTCGTATAG | TbGT8 pSY45_pDS66 |
| | SMD352 | catcgaaactacaCGCAGCCAGGATCCATAACTTC | |
| | SMD353 | atcctggctgcgTGTAGTTTCGATGCCGTTTTCG | TbGT8 3’ homologous flank |
| | SMD354 | gcaggtcgactctagaggatccccgTGCCTTCTACAAGTACGGTTATAC | |
| | SMD355 | gatcGGCCGGCCATGGTTGGACAAATTTTGAGTAGGAGGGGAC | TbGT8 CDS |
| | SMD356 | gatcAGATCTTCACCGCTTGCCGCATGTTG | |
| BSDr TbGT8 KO construct with regulatory elements | SMD400 | aacgacggccagtgaattcgagctcCTTGCATCGCGACTGTTTTTC | 5' TbGT8 homologous flank |
| | SMD401 | aacgagaagaaggGCTGGGGTCCTTTCTTTCCTTG | |
| | SMD402 | aaggaccccagcCCTTCTTCTCGTTAAATGTAC | BSDr with regulatory elements from pNAT |
| | SMD403 | gaaactacatcaAATACTGCATAGATAACAAACG | |
| | SMD404 | tctatgcagtattTGATGTAGTTTCGATGCCGTTTTC | 3' TbGT8 homologous flank |
| | SMD405 | gcaggtcgactctagaggatccccgTGCCTTCTACAAGTACGGTTATAC | |
| RT-qPCR primers for TfR analysis | SMD446 | AACAGATACTGACGGTGTATTGG | ESAG6 |
| | SMD447 | CCACCCTCAACGTACATACTTC | |
| | SMD448 | GACGGAGGTTTGCTGAAAGATA | ESAG7 |
| | SMD449 | GTGAGAACTGACATCACCGTATT | |
| | SMD450 | CGATGATGTGCGAGGTTGT | RBP5 (Tb927.11.12100) |
| | SMD451 | CGTCTGGAACCACTTTCCG | |
| | SMD345 | GACATGTTTGTGCACACCCC | TbGT8 (Tb927.10.12290) |
| | SMD346 | GTAAAGACCCCCAGTTGCGA | |
| | SMD398 | TTCCGCACCCTGAAACTGA | beta-Tubulin |
| | SMD399 | TGACGCCGGACACAACAG | |
| TbGT8 primers binding outwith integration site | SMD357 | TTCCGCTCGTCACTACTGATG | TbGT8 locus 5'F |
| | SMD358 | AAGCCCTCACCTACTAATCCCAC | TbGT8 locus 3' R |

**S1 Table: list of oligonucleotide primers used in this study**
